# Noc corrals migration of FtsZ protofilaments during cytokinesis in *Bacillus subtilis*

**DOI:** 10.1101/2020.08.27.271361

**Authors:** Yuanchen Yu, Jinsheng Zhou, Fredy Gueiros-Filho, Daniel B. Kearns, Stephen C. Jacobson

## Abstract

Bacteria that divide by binary fission form FtsZ rings at the geometric midpoint of the cell between the bulk of the replicated nucleoids. In *B. subtilis*, the DNA- and membrane-binding Noc protein is thought to mediate nucleoid occlusion to prevent FtsZ rings from forming over the chromosome. To explore the role of Noc, we used time-lapse fluorescence microscopy to monitor FtsZ and the nucleoid of cells growing in microfluidic channels. Our data show that Noc does not prevent FtsZ formation over the chromosome or control cell division site selection. Instead, Noc inhibits migration of FtsZ protofilaments from one FtsZ structure to the next. Moreover, we show that FtsZ protofilaments travel due to a local reduction in ZapA association, and the Noc mutant phenotype can be suppressed by ZapA overexpression. Thus, Noc maintains a high local concentration of FtsZ to stabilize FtsZ rings during cytokinesis.

**IMPORTANCE:** In bacteria, a condensed structure of FtsZ (Z-ring) recruits cell division machinery, and Z-ring formation is inhibited over the chromosome by a poorly understood phenomenon called nucleoid occlusion. In *B. subtilis*, nucleoid occlusion has been reported to be mediated by the DNA-membrane bridging protein, Noc. Using time-lapse fluorescence microscopy of cells growing in microchannels, we show that Noc neither protects the chromosome from proximal Z-ring formation nor determines the future site of cell division. Rather, Noc plays a corralling role by preventing protofilaments from leaving a Z-ring undergoing cytokinesis and traveling over the nucleoid.

## INTRODUCTION

Many bacteria grow and divide by a process called binary fission, in which cells increase in biomass and divide into two apparently identical siblings. Prior to division, chromosomes compacted into structures called nucleoids are replicated and segregated to become evenly distributed in the cell (1–4). Next, the early cell division scaffolding protein FtsZ forms an intense ring-like structure in a space between the replicated nucleoids and recruits the machinery for cell envelope biosynthesis (5–8). FtsZ is thought to become concentrated in the interchromosome space, at least in part, by a process called nucleoid occlusion. Nucleoid occlusion is the idea that the presence of the nucleoid inhibits the formation of FtsZ at places along the cell length where the concentration of the chromosomal DNA is greatest (9–12). In so doing, the nucleoid helps guide the division machinery between the replicated chromosomes such that subsequent division will not only perform cytokinesis but ensures each daughter receives one copy of the chromosome (13, 14). The molecular mechanisms underpinning the phenomenon of nucleoid occlusion are poorly understood.

In the Gram positive firmicute *Bacillus subtilis*, nucleoid occlusion is believed to be mediated by the protein, Noc. The *noc* gene was discovered serendipitously when double mutants with the *min* system caused a temperature sensitive defect in cell division (15). Mutation of *noc* alone did not affect growth but resulted in spiral-like intermediates of FtsZ that were observed over the nucleoid, and treatment of a *noc* mutant with a DNA damaging agent resulted in a lethal bisection of trapped chromosomes (15–17). Noc is a ParB-like DNA binding protein that binds to many sites across the chromosome and has an amphipathic helix that promotes membrane association (18, 19). Noc inhibition of FtsZ ring formation requires both DNA and membrane binding activity (18, 19). Thus, Noc may bring regions of the chromosome to the membrane and, thereby, sterically exclude the formation of FtsZ structures around the chromosome.

Here, we monitor FtsZ dynamics in the presence and absence of Noc with chemostatic growth in microfluidic channels. Our data suggest that rather than functioning as an inhibitor of Z-ring formation, Noc plays a stabilizing role on properly assembled FtsZ rings. We observed that mutation of Noc resulted in decondensed spiral-like intermediates of FtsZ over nucleoids, consistent with previous reports. However, the decondensed spiral structures did not appear to form de novo, but rather, split from pre-existing FtsZ rings and migrated to the future site of cell division. We further show that the spiral-like intermediates are locally depleted for association with the Z-ring stabilizing protein ZapA and that artificial overexpression of ZapA could recondense FtsZ in the absence of Noc. Thus, Noc appears to corral the FtsZ ring by preventing the redistribution of decondensed protofilaments in a manner analogous to the roles of septin and in eukaryotic cytokinesis (20–23). Modifying existing models, we speculate that Noc may stabilize FtsZ at least in part by recruiting the nucleoid to the membrane and inhibiting FtsZ-ring migration where chromosome concentration is the highest.

## MATERIALS AND METHODS

### Strains and growth conditions

*Bacillus subtilis* cells were grown in lysogeny broth (LB) (10 g tryptone, 5 g yeast extract, 10 g NaCl per L) or on LB plates fortified with 1.5% Bacto agar at 37 °C. Antibiotics were added when needed at the following concentrations: 100 μg/ml spectinomycin, 5 μg/ml kanamycin, 10 μg/ml tetracycline, 5 μg/ml chloramphenicol, mls (macrolide-lincosamide-streptogramin B, 1 μg/ml erythromycin, 25 μg/ml lincomycin). For *Physpank* promoter dependent gene expression, 1 mM isopropyl-β-d-1-thiogalactopyranoside (IPTG) was added to the medium, if not otherwise indicated.

### Strain construction

All strains were generated by direct transformation of the *B. subtilis* ancestral strain 3610 with enhanced frequency of natural competence for DNA uptake, DK1042 or transduced into DK1042 by SPP1 mediated transduction (24, 25). Cells were mutated for the production of extracellular polysaccharide (EPS) to prevent biofilm formation within the microfluidic device by in-frame markerless deletion of *epsH* genes, encoding enzymes essential for EPS biosynthesis (26, 27). The mNeongreen fluorescent fusion to FtsZ or ZapA, a generous gift of Ethan Garner (Harvard University), was crossed into the indicated genetic background by SPP1 phage mediated transduction, and the antibiotic resistance cassette was eliminated (28). The HBsu-mCherry fusion (4, 29) was transduced into the indicated genetic backgrounds by SPP1 phage mediated transduction.

#### Noc::kan

The *noc::kan* insertion deletion allele was generated with a modified “Gibson” isothermal assembly protocol (30). Briefly, the region upstream of *noc* was PCR amplified with the primer pair 3399/3400, and the region downstream of *noc* was PCR amplified with the primer pair 3401/3402. DNA containing a kanamycin resistance gene (pDG780, 31) was amplified with universal primers 3250/3251. The three DNA fragments were combined at equimolar amounts to a total volume of 5 μL and added to a 15 μl aliquot of prepared master mix (see below). The reaction was incubated for 60 min at 50 °C. The completed reaction was then PCR amplified with primers 3399/3402 to amplify the assembled product. The amplified product was transformed into competent cells of PY79 and then transferred to the 3610 background with SPP1-mediated generalized transduction. Insertions were verified by PCR amplification with primers 3399/3402.

A 5X isothermal assembly reaction buffer (500 mM Tris-HCL pH 7.5, 50 mM MgCl2, 50 mM DTT (Bio-Rad), 31.25 mM PEG-8000 (Fisher Scientific), 5.02 mM NAD (Sigma Aldrich), and 1 mM of each dNTP (New England BioLabs)) was aliquoted and stored at −80° C. An assembly master mixture was made by combining prepared 5X isothermal assembly reaction buffer (131 mM Tris-HCl, 13.1 mM MgCl2, 13.1 mM DTT, 8.21 mM PEG-8000, 1.32 mM NAD, and 0.26 mM each dNTP) with Phusion DNA polymerase (New England BioLabs; 0.033 units/μL), T5 exonuclease diluted 1:5 with 5X reaction buffer (New England BioLabs; 0.01 units/μL), Taq DNA ligase (New England BioLabs; 5328 units/μL), and additional dNTPs (267 μM). The master mix was aliquoted as 15 μL and stored at −80 °C.

#### amyE::Physpank-mRFPmars

To generate the IPTG inducible construct for mRFPmars expression, the gene encoding mRFPmars was PCR amplified from pmRFPmars (Addgene) with primers 4572/4573. The PCR product was purified, digested with NheI and SphI, and cloned into the NheI and SphI sites of pDR111 containing the *Physpank* promoter, the gene conducing the LacI repressor protein, and an antibiotic resistance cassette between the arms of the *amyE* gene (generous gift from David Rudner, Harvard Medical School) to generate pDP430. The pDP430 plasmid was transformed into DK1042 to create DK3391.

#### ycgO::PsigA-mRFPmars

The constitutive mRFPmars construct was generated by isothermal assembly and direct transformation of the linear fragment into *B. subtilis.* The fragments upstream and downstream of *ycgO* were PCR amplified with primers 5270/5271 and 5272/5273, respectively, with chromosomal DNA from strain DK1042 as a template. The fragment containing the *P_sigA_* promoter was PCR amplified with primers 5268/5269 and chromosomal DNA from strain DK1042 as a template. The fragment containing mRFPmars and the spectinomycin resistance cassette was PCR amplified with primers 5266/5267 and chromosomal DNA from strain DK3391 as a template. The four PCR products were purified and mixed in an isothermal assembly reaction that was subsequently amplified by PCR with primers 5270/5273.

### Microfluidic system

The microfluidic device was fabricated through a combination of electron-beam lithography, contact photolithography, and polymer casting (32). Briefly, the microfluidic device is comprised of fluid and control layers both cast in poly(dimethylsiloxane) (PDMS) and a glass coverslip. The fluid layer lies between the control layer and glass coverslip and contains the microchannel array to trap the bacteria. Media and cells are pumped through the microfluidic channels by on-chip valves and peristaltic pumps that are controlled pneumatically through the top control layer. Each pneumatic valve is controlled by software to apply either vacuum (0.3 bar) or pressure (1.3 bar) to open or close individual valves, respectively. The device design is shown in Figure S1.

### On-device cell culture

Prior to loading cells into the microfluidic device, the microchannels were coated with 1% bovine serum albumin (BSA) in LB for 1 h to act as a passivation layer. Then, all the channels were filled with LB media containing 0.1% BSA. A saturated culture of cells (~10 μL) was added through the cell reservoir and pumped into the cell-trapping region. During cell loading, vacuum was applied to the control layer to lift up the microchannel array. After a sufficient number of cells were pumped underneath the channel array, positive pressure was applied to trap individual cells in those channels. Media was pumped through the microchannels to flush away excess cells and maintain steady-state cell growth in the channel array.

### Time-lapse image acquisition

After inoculation in the microfluidic channels, a period of roughly 3 h elapsed during which cells adjusted to the growth conditions, and steady-state cell growth was maintained and monitored over the next 21 h. Fluorescence microscopy was performed either on a Nikon Eclipse Ti-E microscope or an Olympus IX83 microscope. The Nikon Eclipse Ti-E microscope was equipped with a 100x Plan Apo lambda, phase contrast, 1.45 N.A., oil immersion objective and a Photometrics Prime95B sCMOS camera with Nikon Elements software (Nikon, Inc.). Fluorescence signals from mCherry and mNeongreen were captured from a Lumencor SpectraX light engine with matched mCherry and YFP filter sets, respectively, from Chroma. The Olympus IX83 microscope was equipped with an Olympus UApo N 100x/1.49 Oil objective and a Hamamatsu EM-CCD Digital Camera operated with MetaMorph Advanced software. Fluorescence signals from mCherry, mRFPmars, and mNeongreen were excited with an Olympus U-HGLGPS fluorescence light source with matched TRITC, TRITC and FITC filters, respectively, from Semrock. Images were captured from at least eight fields of view at 2 min intervals. The channel array was maintained at 37 °C with a TC-1-100s temperature controller (Bioscience Tools). For all direct comparisons, the same microscope and settings were used.

### Data analysis

An adaptation period following exposure to illumination was observed; thus, data analysis was restricted to periods of steady state. Cell identification and tracking were analyzed by MATLAB programs (The MathWorks, Inc.) (32). The program extracted fluorescence intensity along a line profile down the longitudinal center of each microchannel in the array. The cytoplasmic mCherry line profile showed a flat-topped peak on the line where a cell was located, and a local 20% decrease in fluorescence intensity was used to identify cell boundaries after division. Division events were conservatively measured as the time at which one cell became two according to the decrease in fluorescence intensity. Moreover, cell bodies were tracked from frame to frame in order to construct lineages of cell division, and cell body intensity was determined by the integration of the cytoplasmic mCherry signal intensity within the confines of the cell. Signals from HBsu-mCherry and mNeongreen-FtsZ were similarly tracked and measured along the length of the cell. FtsZ line profiles were normalized by cell body intensity in order to minimize intensity differences among frames and across different fields of view.

### Microscopic images acquisition on agarose pads

Cells were grown to mid-log phase in liquid culture and then imaged on 1% agarose pads in S750 media (33). Fluorescence microscopy was performed with a Nikon 80i microscope with a Nikon Plan Apo 100X phase contrast objective and an Excite 120 metal halide lamp. Cytoplasmic mRFPmars was visualized with a C-FL HYQ Texas Red filter cube. mNeongreen-FtsZ was visualized with a C-FL HYQ FITC filter cube. Images were captured with a Photometrics Coolsnap HQ2 camera in black and white, false colored, and superimposed with Metamorph image software.

**Table 1:** Strains

| Strain | Genotype |
| --- | --- |
| DK1042 | Wild type |
| DK3391 | *amyE::P<sub>hyspank</sub>-mRFPmars spec* |
| DK5133 | *mNeongreen-ftsZ ΔepsH amyE::P<sub>hyspank</sub>-mcherry spec* |
| DK5712 | *mNeongreen-ftsZ ΔepsH sacA::hbsu-mCherry cat* |
| DK5820 | *mNeongreen-ftsZ ΔepsH amyE::P<sub>hyspank</sub>-mcherry spec noc::kan* |
| DK6372 | *mNeongreen-ftsZ ΔepsH sacA::hbsu-mCherry cat noc::kan* |
| DK8138 | *zapA-mNeongreen ΔepsH amyE::P<sub>hyspank</sub>-mcherry spec* |
| DK8139 | *mNeongreen-ftsZ ΔepsH amyE::P<sub>hyspank</sub>-zapA spec ycgO::P<sub>sigA</sub>-mRFP mars mls* |
| DK8144 | *mNeongreen-ftsZ ΔepsH amyE::P<sub>hyspank</sub>-zapA spec ycgO::P<sub>sigA</sub>-mRFP mars mls noc::kan* |
| DK8162 | *mNeongreen-ftsZ ΔepsH ycgO::P<sub>sigA</sub>-mRFP mars mls* |
| DK8170 | *mNeongreen-ftsZ ΔepsH ycgO::P<sub>sigA</sub>-mRFP mars mls noc::kan* |
| DK8172 | *zapA-mNeongreen ΔepsH amyE::P<sub>hyspank</sub>-mcherry spec noc::kan* |

**Table 2:**
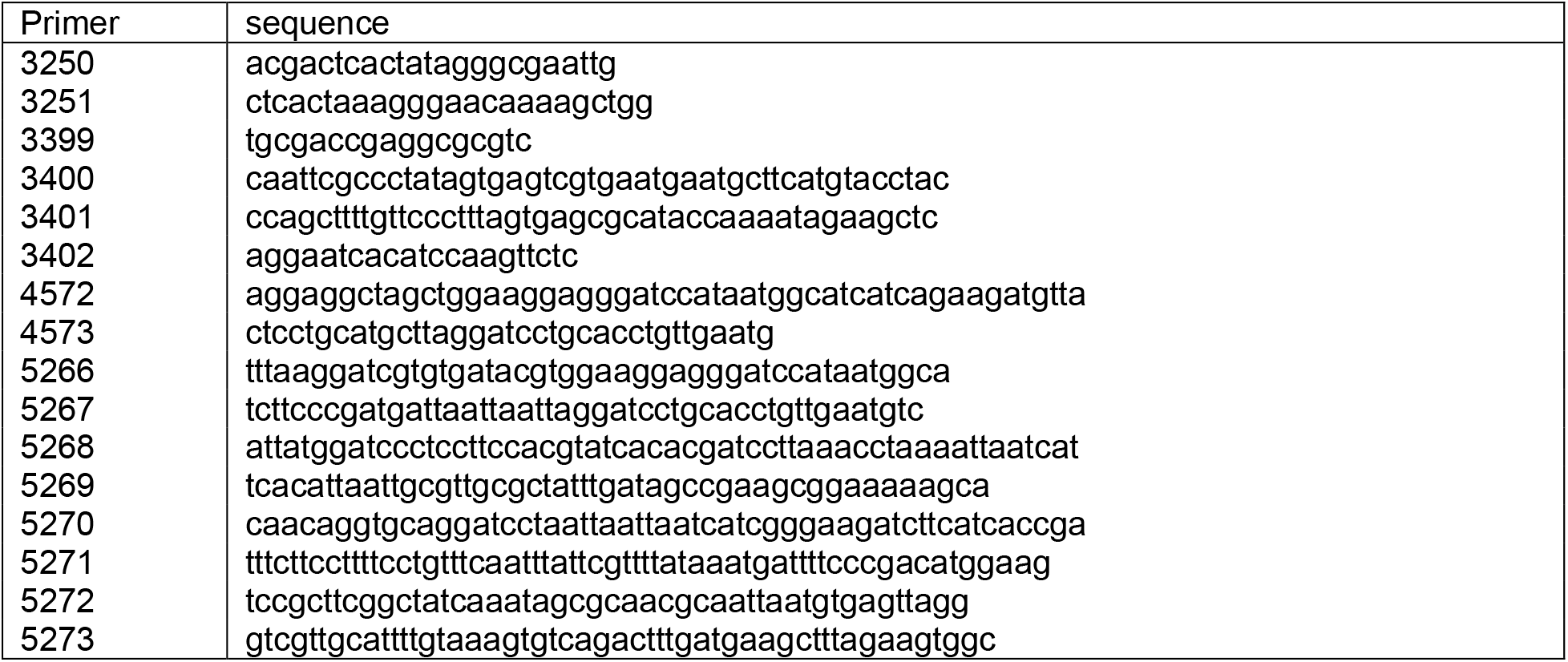
Primers

## RESULTS

### Noc prevents FtsZ from travelling over the chromosome

In *B. subtilis*, the protein Noc is thought to function in nucleoid occlusion by preventing FtsZ formation and subsequent cell division from occurring over the chromosome. We used a microfluidic platform with microchannel arrays to quantitatively study the role of Noc on bacterial cell division during chemostatic growth. Fluorescence images of cells expressing cytoplasmic red fluorescent protein were captured every 2 min after cells entered steady state growth, and cell division was defined as a 20% decrease in fluorescence intensity along the longitudinal axis of the cell (**Figure 1A**). Consistent with previous reports, microfluidic analysis showed that most of the cell division events occurred in the middle of the *noc* mutant (**Figure 1B**), with an average division time similar to that of the wild type (**Figure 2A**). The growth rate measured by optical density in liquid culture (**Figure 2B**) and cellular elongation rate per cell length (**Figure 2C**) of the noc mutant were similar to that of wild type. We conclude that disruption of *noc* had minimal effects on the growth and division of the cell.

**Figure 1:**
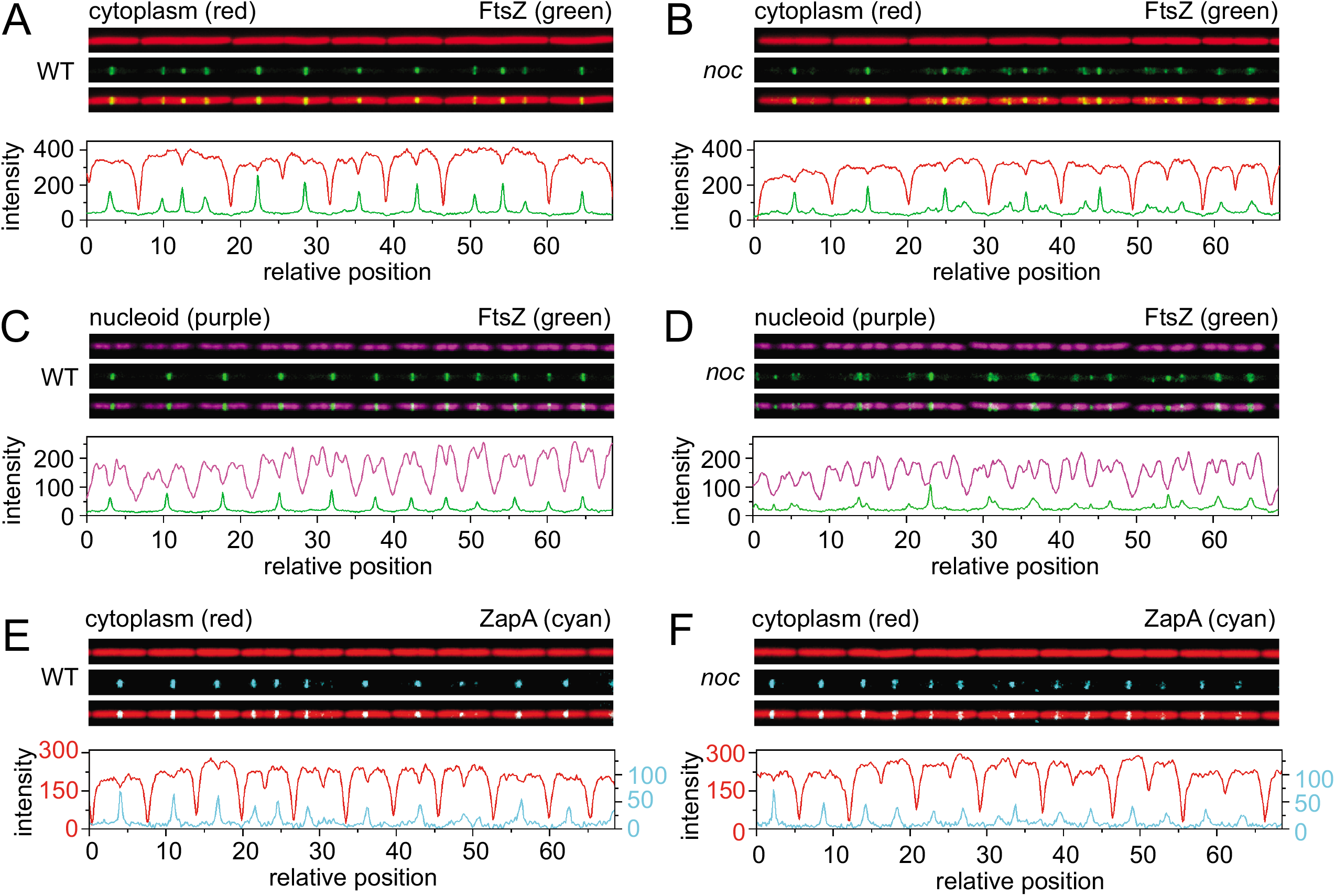
Microfluidic analysis of growth, division, and chromosome segregation in wild type and *noc* mutants. Fluorescence microscopy of wild type (A, C, E) and *noc* mutants (B, D, F) growing at steady state in a microfluidic channel. A and B) Fluorescence microscopy of wild type (DK5133, A) and the *noc* mutant (DK5820, B). Cytoplasmic mCherry false colored red (top), mNeongreen-FtsZ false colored green (middle), and an overlay of the two colors (bottom). Graphs are a quantitative analysis of mCherry fluorescence intensity (red line) and mNeongreen fluorescence intensity (green line) to match the fluorescence images immediately above. C and D) Fluorescence microscopy of wild type (DK5712, C) and a *noc* mutant (DK6372, D). Chromosomes (HBsu-mCherry) false colored purple (top), mNeongreen-FtsZ false colored green (middle), and an overlay of the two colors (bottom). Graphs are a quantitative analysis of mCherry fluorescence intensity (purple) and mNeongreen fluorescence intensity (green) to match the fluorescence images immediately above. E and F) Fluorescence microscopy of wild type (DK 8138, E) and a *noc* mutant (DK8172, F). Cytoplasmic mCherry false colored red (top), ZapA-mNeongreen false colored cyan (middle), and an overlay of the two colors (bottom). Graphs are a quantitative analysis of mCherry fluorescence intensity (red) and mNeongreen fluorescence intensity (cyan) to match the fluorescence images immediately above. For panels E and F) two different Y-axes are used due to lower intensity fluorescence from the ZapA-mNeongreen construct. The left axis corresponds to the mCherry signal (red) and the right axis corresponds to the ZapA-mNeongreen signal (cyan). All images are reproduced at the same magnification.

**Figure 2:**
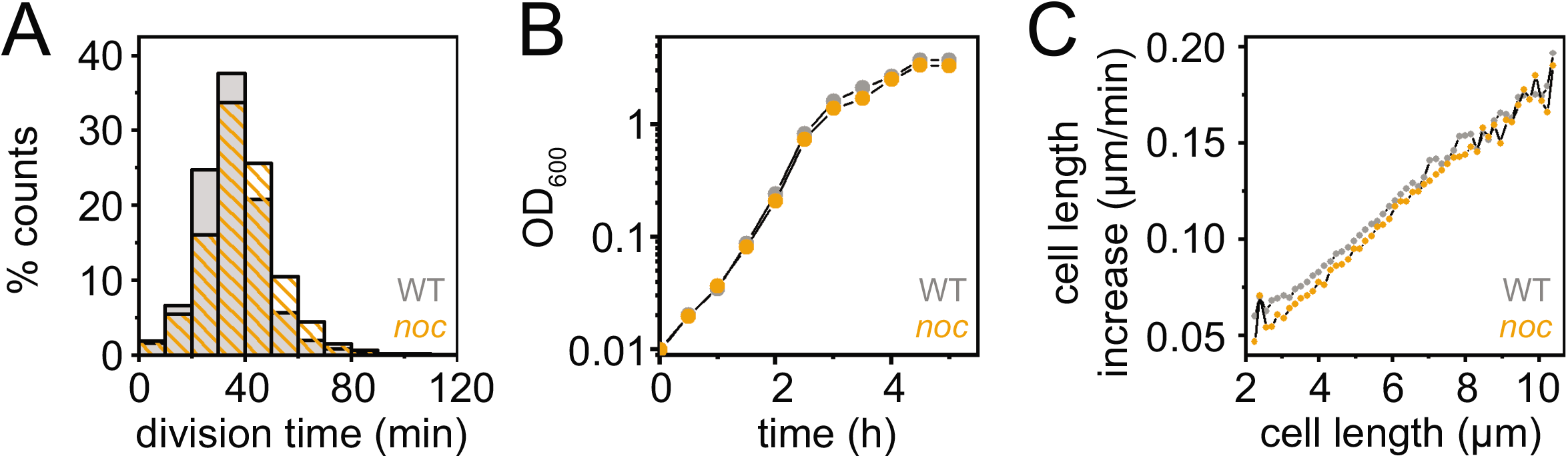
Cells mutated for Noc grow like wild type. A) Histogram of the division time of individual cells of wild type (DK5133, gray) and a *noc* mutant (DK5820, orange) measured by microscopic analysis. Division events were defined by a local 20% decrease in mCherry (cytoplasmic) fluorescence intensity below a threshold value, and the division time was the time between two consecutive division events. More than 2800 division events were counted per dataset. B) Growth curve of wild type (gray) and a *noc* mutant (orange) growing in highly agitated LB broth at 37 °C. Optical density was measured with a spectrophotometer at 600 nm wavelength. C) Increase in cell length for wild type (gray) and the *noc* mutant (orange) was measured as the rate at which the cell poles moved apart from one another relative to the current cell length.

Nucleoid occlusion is thought to be mediated by preventing FtsZ ring formation in the vicinity of the chromosomes. To investigate the localization of the Z-ring, mNeongreen was fused to the N-terminus of FtsZ at the native site in the chromosome, and FtsZ dynamics were tracked in the wild type (**Figure 1A, Movie S1**) and the *noc* mutant (**Figure 1B, Movie S2**). In wild type, FtsZ primarily localized to the midcell and cell poles (**Figure 3A**), but in the *noc* mutant, additional peaks were observed at the one-quarter and three-quarter cell positions (**Figure 3B**). Moreover, the FtsZ foci observed in the *noc* mutant appeared wider and more diffuse (**Figure 1B**). Kymograph analysis showed that in wild type, FtsZ rings appeared as intense bands that disappeared from the cell pole after division and reappeared at the nascent midcell (**Figure 4A**). In the *noc* mutant, however, a subpopulation of FtsZ appeared to split from the primary focus and migrate toward the next midcell as a spiral-like intermediate (**Figure 4B**). We infer that FtsZ ring migration in the *noc* mutant was responsible for the increase in peak intensity observed at positions other than the midcell and poles. We conclude that Noc does not prevent formation of Z-rings over the nucleoid. Rather, Noc stabilizes FtsZ rings by inhibiting a portion of FtsZ from migrating along the length of the cell, likely over the top of the chromosome.

**Figure 3:**
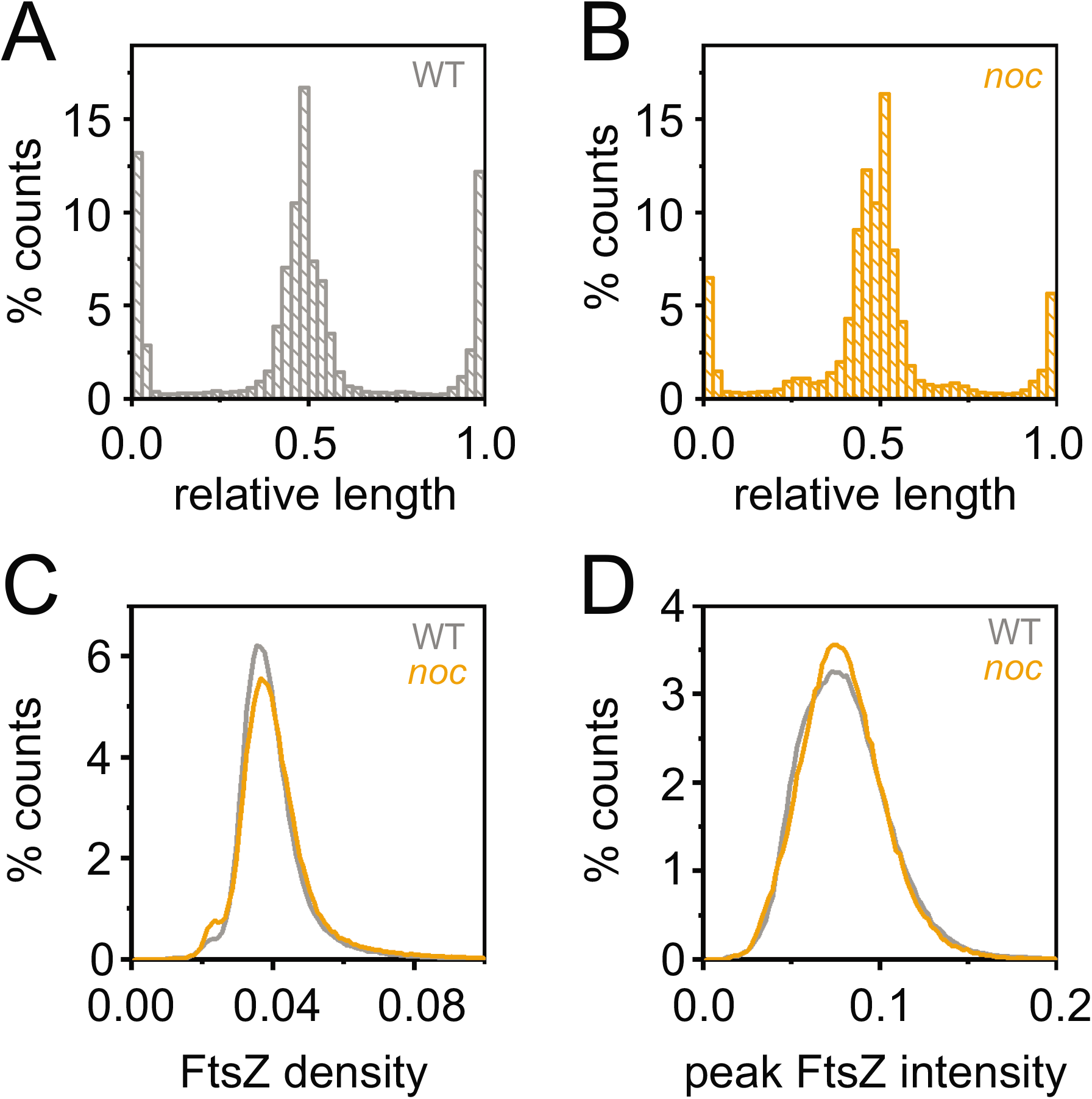
FtsZ intensity at locations other than the poles and midcell increased in the *noc* mutant. A) Histogram of the FtsZ intensity along the relative cell length of wild type (gray). Relative lengths are 0 and 1.0 for the cell poles and 0.5 for the midcell. B) Histogram of the FtsZ intensity along the relative cell length of the *noc* mutant (orange). C) Histogram of FtsZ density in individual cells measured by dividing the total FtsZ fluorescence intensity (Supplemental Fig S2A) by cell length (Supplemental Fig S2B) for wild type (gray) and *noc* mutant (orange). D) Histogram of peak FtsZ fluorescence intensity in individual cells for wild type (gray) and *noc* mutant (orange). Data for each panel were generated by a line scan along the longitudinal axis of the cell to determine the location of fluorescence intensity for each frame during time-lapse microscopy. Data were taken from 13000 wild type (DK5133) for > 420,000 measurements and 5000 *noc* mutant (DK5820) for > 170,000 measurements.

**Figure 4.**
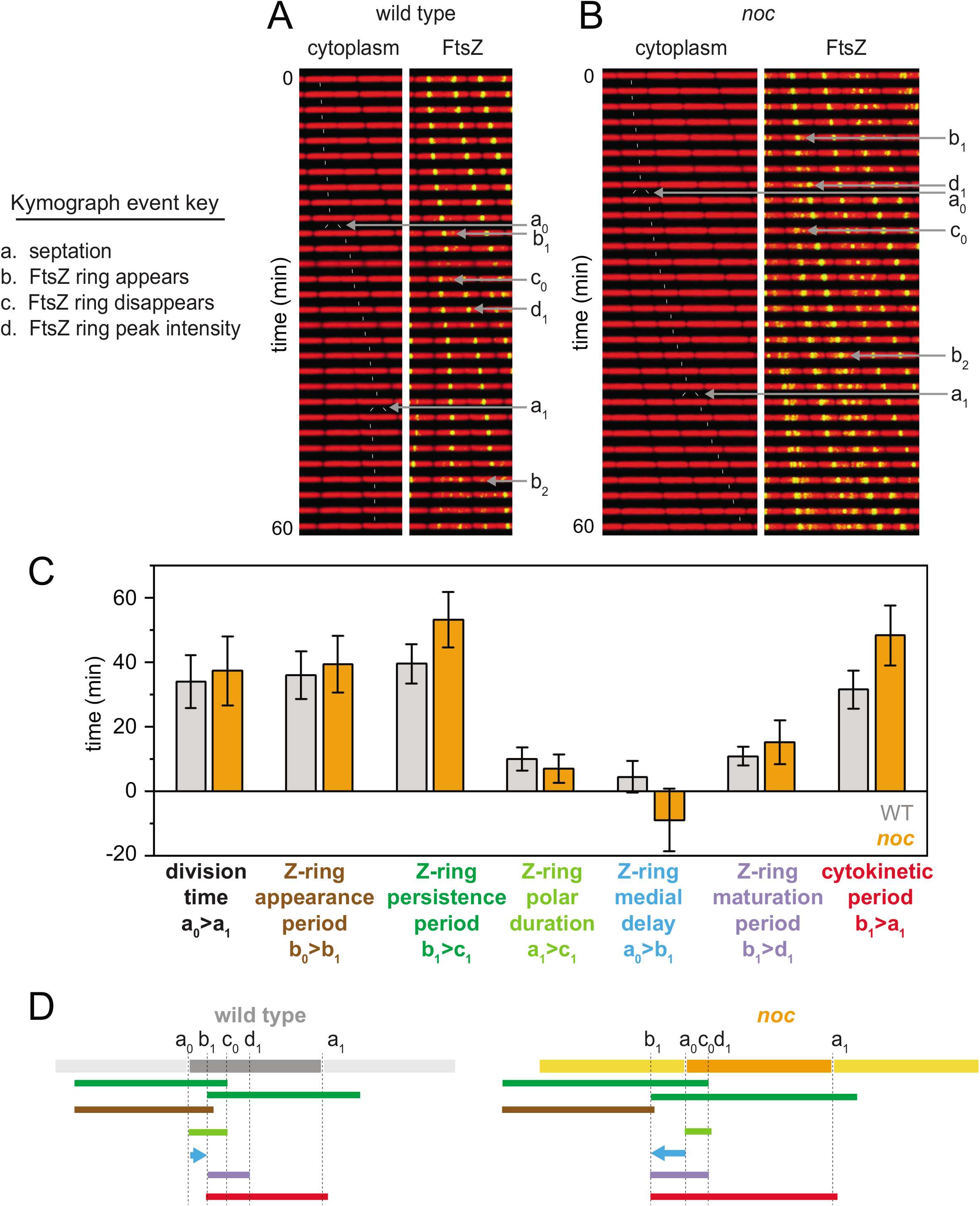
In *noc* mutants, a subset of FtsZ disassociates from the Z-ring and travels to the future site of cell division, which causes the Z-ring medial delay time to become negative. A and B) Kymograph analysis of wild type (DK5133, A) and *noc* mutant (DK5820, B) grown in a microfluidic channel and imaged every 2 min. Cytoplasmic mCherry signal is false colored red (left) and overlaid with mNeongreen-FtsZ that is false colored green (right). Events necessary for defining division parameters are indicated and labeled a, septation; b, appearance of a nascent Z-ring; c, disappearance of a Z-ring; d, FtsZ peak intensity achieved. Each event designation is given a number: 0 for the preceding generation, 1 for the current generation, and 2 for the subsequent generation. Thin white lines are included to indicate cell tracking and lineage analysis. C) Graphs of 100 manually tracked cell division cycles for wild type (gray) and *noc* mutant (orange) presented as bars of average values and whiskers of standard deviation for the following parameters: “division time” is the time between septation events (between consecutive “a” events); “Z-ring appearance period” is the time between the appearance of one Z-ring and another (between consecutive “b” events); “Z-ring persistence period” is the time between the appearance of a Z-ring and the disappearance of that Z-ring (between consecutive “b” and “c” events); “Z-ring polar duration” is the time between a septation event and the disappearance of the Z-ring resulting from that septation event (between consecutive “a” and “c” events); “Z-ring medial delay” is the time between a septation event and the appearance of a Z-ring that will eventually give rise to the next medial division event (between an “a” event and a “b” event that will give rise to the next round of septation); “Z-ring maturation period” is the time between Z-ring appearance and when that Z-ring achieves peak local intensity (between consecutive “a” and “d” events); and “cytokinetic period” is the time between Z-ring appearance and septation directed by that Z-ring (between a “b” event and the “a” event that is caused by that particular Z-ring). In the case of the *noc* mutant, the Z-ring that will form the future division site first appears by splitting from the pre-existent Z-ring, and the appearance period is defined as the time at which the Z-ring visibly splits from its progenitor. Note that the Z-ring medial delay of the *noc* mutant was negative on average because the medial Z-ring that would eventually promote cell division was formed in the preceding generation. Histograms for each bar are presented in Supplemental Fig S2C-H. D) Timelines of the various events indicated in the bar graph, color coded to match the indicated parameter of like color above, and annotated with relevant events marked by the defining letters.

To determine Z-ring localization relative to the chromosomes, the nucleoid was fluorescently labeled by fusion of a fluorescent protein, mCherry, to the nucleoid binding protein HBsu and incorporated in merodiploidy at an ectopic site (4, 29). During growth of the wild type, nucleoids appeared as intense fluorescence in the cytoplasm and were sometimes observed to form bilobed masses indicative of partial replication and segregation (**Figure 1C, Figure 5A, Movie S3**). Moreover, FtsZ rings formed preferentially between partially replicated nucleoid lobes over the low density inter-chromosome region (**Figure 1C, Figure 5A, Movie S3**). Chromosome structure in the *noc* mutant appeared perturbed in which nucleoids experienced wider and irregular separation, but FtsZ rings still predominantly concentrated between chromosomes (**Figure 1D, Figure 5B, Movie S4**). In the *noc* mutant however, FtsZ structures were also observed over the chromosome mass (**Figure 1D, Figure 5B, Movie S4**). Kymograph analysis indicated that 61% of the cells had diffuse FtsZ foci migrate from the polar ring to the future division site by traveling over the nucleoid, compared to only 8% in the wild type (**Figure 5C**). We conclude that in the *noc* mutant, a portion of FtsZ splits from the primary ring and migrates atop the chromosome to coalesce at the next midcell position. Thus, Noc does not prevent the de novo formation FtsZ foci over the nucleoid but rather corrals FtsZ at its present location.

**Figure 5:**
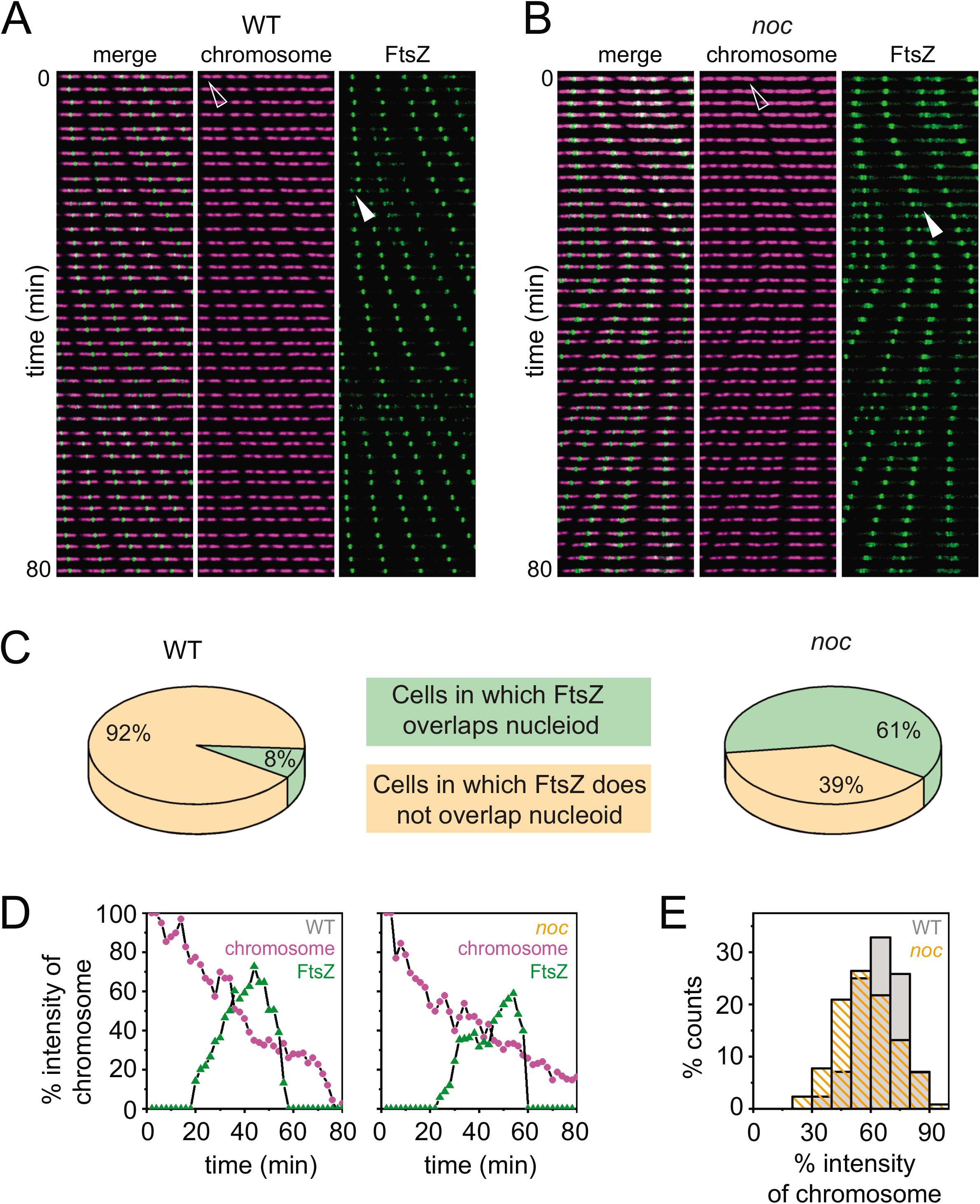
The FtsZ-ring does not form in the chromosome gap earlier in the *noc* mutant. A and B) Kymograph analysis of wild type (DK5712, A) and a *noc* mutant (DK6372, B) grown in a microfluidic channel and imaged every 2 min. Chromosomes (HBsu-mCherry) false colored purple and mNeongreen-FtsZ false colored green. All images reproduced at the same magnification. Left, color overlay; center, HBsu-mCherry; right; mNeongreen-FtsZ. Open caret indicates the nucleoid intensity monitored in the corresponding graphs in panel D. Closed caret indicates the FtsZ intensity monitored in the corresponding graphs in panel D. C) Pie charts indicate the percentage of cells in which FtsZ rings are found to overlap (green) or not overlap (tan) with the bulk of the nucleoid mass in wild type and the *noc* mutant. 200 cells were counted for each background. D) HBsu-mCherry (chromosome, purple) and mNeongreen-FtsZ (FtsZ, green) intensity at mid-chromosome over time. Both values were normalized by the maximum intensity of the nucleoid to minimize intensity differential across different time points. Left panel shows a representative result of a single wild type cell (carets in panel A), and right panel shows a representative result of a single *noc* mutant cell (carets in panel B). E) Histogram of the chromosome intensity at which FtsZ first localized precisely at the newly formed chromosome gap. Data are taken from 200 cells, similar to data in panel D.

### Noc corralling of the Z-ring alters dynamics but has a neutral outcome on division

Overexpression of FtsZ is reported to result in decondensed FtsZ structures that travel in a manner similar to those observed in the *noc* mutant (34–37). Thus, one way in which the *noc* mutant could give rise to traveling foci is if it exhibited an increase in FtsZ expression. To measure the amount of FtsZ, fluorescence intensity was measured per cell in the wild type and *noc* mutant. The total amount of FtsZ appeared slightly higher in the *noc* mutant than the wild type (**Figure S2A**), but on average, the *noc* cells were also slightly longer (**Figure S2B**). Consequently, the FtsZ intensity per unit length was the same in both cell types (**Figure 3C**). Furthermore, both the wild type and the *noc* mutant showed a similar maximum FtsZ peak intensity (**Figure 3D**). We conclude that, consistent with previous reports, the absence of the Noc protein did not affect the FtsZ concentration nor did its absence alter the ability of FtsZ rings to reach peak local intensity. We infer that absence of Noc leads to alterations in Z-ring function and dynamics.

To explore Z-ring dynamics, 100 cells each were chosen at random from the wild type and *noc* mutant, and several parameters were manually quantified (38). Four events were sequentially measured: the appearance of an FtsZ ring, accumulation of FtsZ to peak intensity, cell division, and disappearance of the FtsZ ring. Four of the seven parameters were similar for both cell types, and three were different (**Figure 4C, Figure S2C-H**). For example, the FtsZ persistence time, defined as the time between the appearance and disappearance of an FtsZ ring, was longer in the *noc* mutant (**Figure 4C, 4D, Figure S2D**). The longer persistence time was not due to decreased rate of polar Z-ring disassembly (**Figure 4C, 4D, Figure S2E**), as had been observed with cells mutated for the Min system (38). Instead, the medial delay time, defined as the time between cell division and appearance of the next Z-ring, was negative as future rings were formed by the splitting from a Z-ring soon after coalescence (**Figure 4C, 4D, Figure S2F**). Combined, the *noc* mutant also experienced a net increase in the cytokinetic period, defined as the time between Z-ring appearance and cell division mediated by that Z-ring (**Figure 4C, 4D, Figure S2H**), largely due to the reduction in the medial delay. Thus, FtsZ forms a ring at the midcell earlier in the noc mutant, but faster formation does not result in faster division, likely because cytokinesis is limited by the recycling of other divisome components from the previous site of division.

The negative medial delay time in the *noc* mutant might suggest that FtsZ finds the inter-chromosome region and future site of cell division earlier compared to wild type. To investigate this hypothesis, we tracked chromosomes and FtsZ foci simultaneously. We defined the chromosome segregation time as the time needed for one chromosome density to become two chromosomes, as measured by a local 40% decrease in HBsu-mCherry fluorescence intensity in the inter-chromosome region. The chromosome segregation time was the same in both the wild type and *noc* mutant indicating that Noc did not have a major role in either genome replication or separation (**Figure S2I**). In wild type, an FtsZ focus appeared in the inter-chromosome region when the remaining chromosome intensity dropped to 64 ± 11%, and the FtsZ focus appeared in the inter-chromosome region in the *noc* mutant at approximately the same chromosome intensity (**Figure 5D,E**). Thus, in the absence of Noc, the decondensed FtsZ travels from a condensed Z-ring over top of the chromosome, but otherwise, forms a new Z-ring at the same position as wild type. Thus, the traveling FtsZ focus appeared to be a neutral alternative to full FtsZ disassembly and its *de novo* synthesis at the midcell.

### ZapA overexpression stabilizes Z-rings in the *noc* mutant

In the *noc* mutant, the FtsZ ring was decondensed and appeared to travel in a helical pattern across the nucleoid. FtsZ polymerizes as protofilaments which, in turn, are condensed into Z-rings by a protofilament bundling protein called ZapA (39–42). FtsZ might appear to be decondensed in the *noc* mutant, if the traveling focus had less ZapA than medial or polar Z-rings. To investigate ZapA localization, mNeongreen was fused to the N-terminus of ZapA at the native site in the chromosome, and dynamics were tracked in the wild type and the *noc* mutant. In wild type, ZapA localized to the midcell and cell poles much like FtsZ (**Figure 1E, Movie S5**). In the *noc* mutant, ZapA also localized to the midcell and cell poles, but unlike FtsZ, decondensed ZapA structures were not observed (**Figure 1F, Movie S6**). Kymograph analysis indicated that the dynamics of ZapA was similar in both wild type and the *noc* mutant (**Figure S3**), and unlike FtsZ, the ZapA foci did not appear to travel over the chromosome in the *noc* mutant (**Figure 4B**). We conclude that in the absence of Noc, FtsZ foci become decondensed and travel over top of the chromosome in a manner correlated with our inability to detect ZapA in the decondensed rings.

If decondensed, traveling FtsZ foci were due to a local reduction in ZapA association, we hypothesized that Z-rings of the *noc* mutant might be recondensed by inducing ZapA overexpression. ZapA expression was controlled by cloning the *zapA* gene under the regulation of the IPTG inducible *Physpank* promoter and inserted as a merodiploid at the ectopic *amyE* site of a *noc* mutant that expressed both cytoplasmic mRFPmars and mNeongreen-FtsZ. In the absence of IPTG, the cells exhibited decondensed Z-rings consistent with the *noc* mutant phenotype (**Figure 6A**). As the amount of IPTG was increased to 0.01 mM, condensation of the Z-rings was observed such that they resembled those of the wild type (**Figure 6A**). At IPTG concentrations above 0.01 mM, however, decondensed Z-rings reappeared, and higher levels of induction abolished cell division altogether (**Figure 6A**). An excess of IPTG also resulted in decondensed Z-rings in the wild type but did not appear to inhibit cell division as severely as when Noc was absent (**Figure 6B**). We infer that the *noc* mutant is sensitized to overexpression of ZapA but that Z-rings could be recondensed by precise titration.

**Figure 6:**
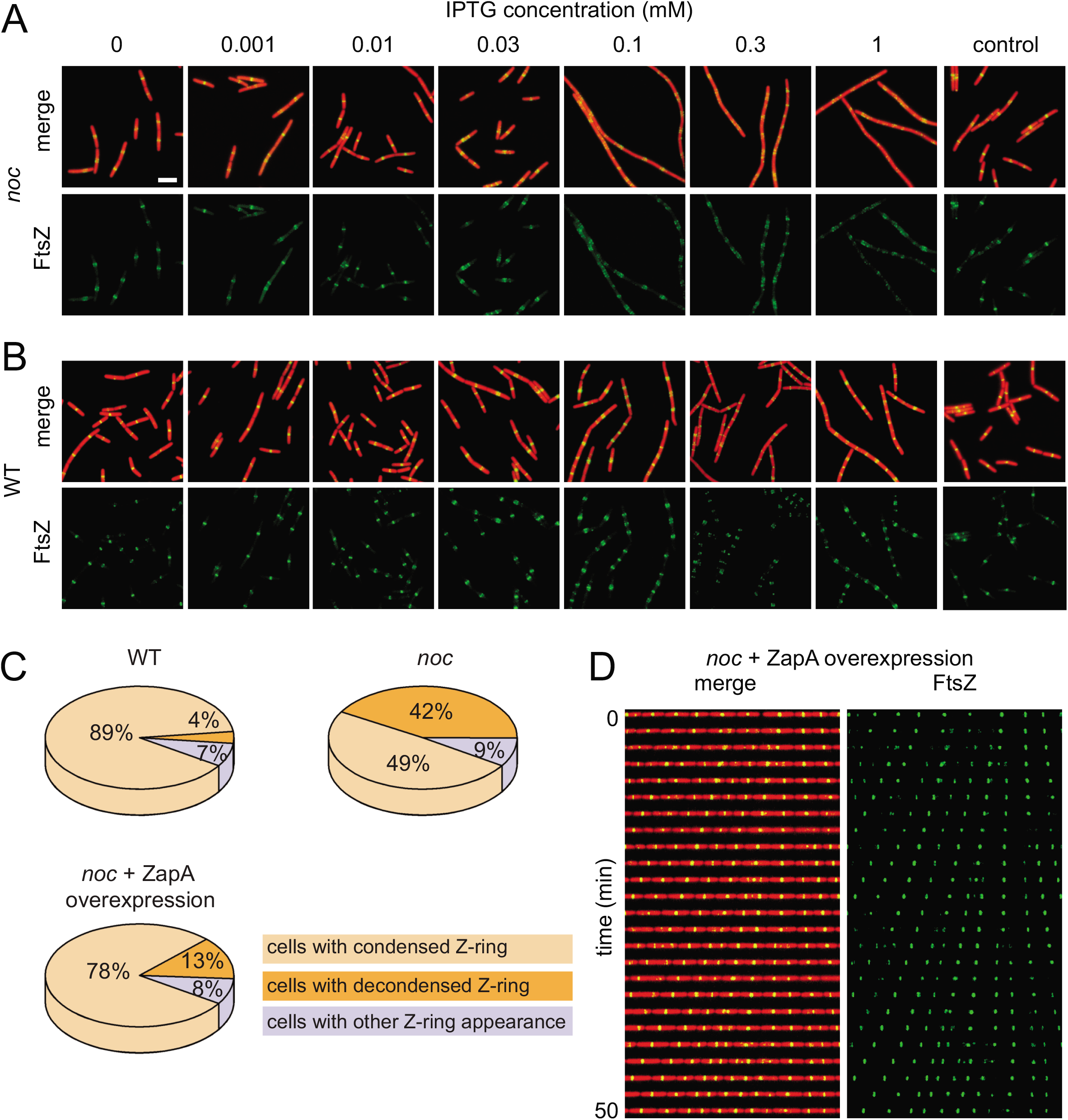
Overexpression of ZapA stabilizes FtsZ-rings in a *noc* mutant. A and B) Fluorescence microscopy of a *noc* mutant (DK8144, A) and wild type (DK8139, B) containing an IPTG-inducible copy of ZapA grown in broth culture containing the indicated amount of IPTG. A *noc* mutant (DK8170) and wild type (DK8162) lacking the inducible ZapA construct were used as controls, respectively. Constitutively expressed cytoplasmic mRFPmars false colored red and mNeongreen-FtsZ false colored green. Scale bar is 5 μm and applies to all panels. C) Pie charts indicate the percentage of cells for each genotype that exhibited condensed Z-rings (tan), decondensed Z-rings (orange), and Z-rings with other patterns (lavendar) in the indicated backgrounds. 200 cells each for wild type (DK8162), *noc* mutant (DK8170), and *noc* mutant in which ZapA was induced with 0.01 mM IPTG (DK8144) were analyzed to generate each graph, respectively. D). Kymograph analysis of a *noc* mutant (DK8144) in which ZapA was induced with 0.01 mM IPTG. Constitutively expressed cytoplasmic mRFPmars false colored red and mNeongreen-FtsZ false colored green. To assemble a kymograph, a single microfluidic channel is imaged at 2 min intervals.

Based on qualitative appearance of condensed Z-rings, we chose 0.01 mM IPTG induction of ZapA as our standard condition for increasing Z-ring condensation in the *noc* mutant. Quantitatively, the FtsZ ring appeared to be condensed at either the midcell or cell pole in 89% of wild type cells and 49% of *noc* mutant cells (**Figure 6C)**. Induction of ZapA in the *noc* mutant with 0.01 mM IPTG, increased the percentage of cells that had a condensed Z-ring to 78% (**Figure 6C**). We conclude that induced ZapA expression was sufficient to recondense the Z-rings in the *noc* mutant. Moreover, kymograph analysis indicated that when ZapA expression was induced in the *noc* mutant, the Z-rings appeared less mobile and perhaps stationary at the midcell and poles (**Figure 6D, Movie S7**). We conclude that overexpression of the Z-ring stabilizing protein ZapA can compensate for the absence of Noc and prevent Z-rings from traveling over the chromosome. We further conclude that the primary function of Noc is not to prevent Z-ring formation over the chromosome per se but rather function, like ZapA, in corralling Z-rings.

## DISCUSSION

One of the primary topological determinants of bacterial cell division is reported to be the phenomenon of nucleoid occlusion, in which FtsZ rings are prevented from forming over the bulk of the chromosome (10, 13, 43). In *B. subtilis*, nucleoid occlusion is thought to be mediated by the Noc protein which, in turn, directly or indirectly inhibits FtsZ polymerization. In support of an inhibitory role, the *noc* gene was originally identified as being synthetically lethal in the absence of the Min system, such that in a *min noc* double mutant, FtsZ polymerized in an unrestricted and unproductive manner throughout the cell (15). Moreover, overexpression of Noc exhibited a modest inhibition of cell division, which could be enhanced in the presence of a high-copy number plasmid containing a Noc-binding sequence (15, 18). Noc bound to many sites across the chromosome except the terminus region, but because the terminus is the part of the nucleoid that remains at midcell after replication, Noc supposedly acts as a timer for midcell Z-ring assembly (18, 44). If and how Noc functions as an inhibitor of FtsZ is unknown. Here, we perform time-lapse video microscopy of FtsZ ring formation in the absence of Noc, and our data suggest that rather than playing an inhibitory role, Noc may function to corral or concentrate FtsZ in the vicinity of a pre-existing Z-ring.

Noc is reported to mediate nucleoid occlusion and direct FtsZ to form an intense pre-divisional focus between the segregated nucleoids at the midcell. Previous work monitoring the first cell division event after spore germination, however, challenged the idea that Noc was involved in division-site selection as FtsZ formed a ring at the midcell between chromosomes with high fidelity, even in the absence of Noc (44). Our time-lapse observations and quantitative microscopy also argue against a role of Noc in cell division-site selection under standard growth conditions. We find that the FtsZ ring formed in the middle of the bilobed nucleoid structure when the amount of chromosome material (indirectly indicated by the presence of the HBsu nucleoid binding protein) was spatially reduced to 60% of maximum intensity in both the wild type and the Noc mutant (7). Thus, if occlusion by the nucleoid directs Z-ring localization, the occlusion somehow becomes abrogated when the local nucleoid volume is reduced by approximately half. By whatever the mechanism this event is interpreted, Noc does not appear to be involved (44).

Noc is thought to guide FtsZ localization by inhibiting the formation of FtsZ over the bulk of the nucleoid. Previous work shows that when Noc is mutated, decondensed helical foci of FtsZ are superimposed over the nucleoid consistent with the idea that Noc prevents formation of FtsZ over the chromosome (15). Although we also observe superposition of decondensed FtsZ structures and the nucleoid, their superposition is not the result of de novo synthesis but rather of directional migration of an FtsZ subpopulation from one midcell Z-ring to the next site of cell division (**Figure 7**). Thus, Noc corrals and concentrates FtsZ at the location of a Z-ring. In addition to a DNA binding domain, Noc encodes an amphipathic helix such that the DNA-Noc nucleoprotein complex is recruited to the membrane (19). Thus, extensive DNA-membrane interactions could form a steric block to mediate the corralling phenomenon and prevent FtsZ migration over the chromosome. We conclude that Noc promotes nucleoid occlusion, not by preventing Z-ring formation, but by inhibiting the migration of FtsZ protofilaments.

**Figure 7.**
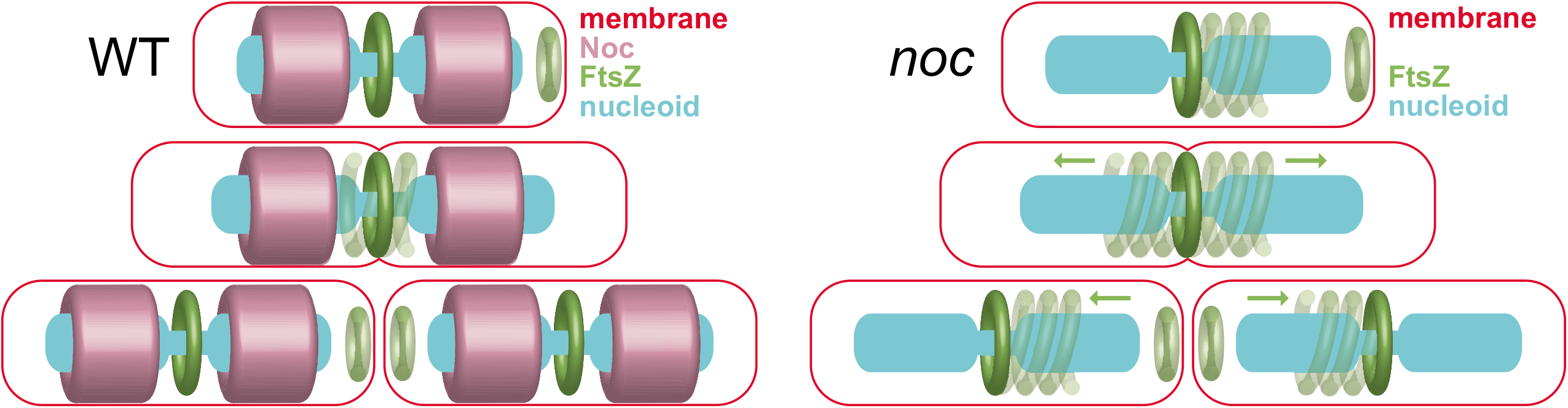
Cartoon model of Noc corralling function. Noc binds to the chromosome and membrane and corrals migration of FtsZ protofilaments during cytokinesis. In the absence of Noc, traveling protofilaments depleted for ZapA travel from a condensed FtsZ focus towards the future site of cell division. FtsZ recondenses at the same position, in the inter-chromosome space where the nucleoid density is reduced regardless of whether Noc is present. Membrane (red), FtsZ (green), Noc (purple), and nucleoid (cyan)

If Noc inhibits the migration of decondensed FtsZ protofilaments, what governs FtsZ condensation in the first place? Traveling FtsZ protofilaments are correlated with an elevated level of FtsZ (45, 46), but FtsZ levels do not change in the absence of Noc. Instead, FtsZ condensation is correlated with the co-localization of the protofilament cross-linking protein ZapA, and the decondensed traveling Z-foci in the *noc* mutant appear to be ZapA-deficient. Consistent with local ZapA depletion, overexpression of ZapA recondenses the FtsZ foci into tight rings, but we infer that this effect is compensatory because the levels of ZapA are, like that observed with FtsZ, unchanged in the *noc* mutant. ZapA may be removed from a subpopulation of FtsZ either stochastically or as a consequence of cytokinesis, and appearance of decondensed FtsZ foci coincides with the start of septum constriction. Recent work suggests that the ZapA-FtsZ complex may oppose constriction, and thus, ZapA binding and removal may be subject to spatio-temporal control (47). Thus, as cytokinesis proceeds, FtsZ protofilaments wander from the active Z-ring, and Noc may use the chromosome as a topological determinant to restrict FtsZ travel.

Our data support a role of Noc in the phenomenon of nucleoid occlusion but not for the purpose of protecting chromosome integrity or directing the location of the future site of cell division. Instead, the DNA-Noc-membrane super-complex sterically hinders treadmilling protofilaments to the cytokinetic ring, after which they are depolymerized by the Min system and repolymerized at the next site cell division. In the *noc* mutant, however, a subpopulation of protofilaments escapes the cytokinetic ring,travels over the chromosome, and arrives at the future site of cell division earlier than the wild type. Nevertheless the cell cycle is largely unperturbed as Z-ring maturation requires a combination of de novo FtsZ synthesis and Min-mediated monomer recycling, and the rest of the division machinery relocates on time. Moreover, overexpression of ZapA compensates for the absence of Noc perhaps by restricting FtsZ migration by increased protofilament crosslinking. We propose a model in which Noc increases division efficiency by forming a barrier that corrals and concentrates cell division machinery. We imagine Noc functions in a manner analogous to eukaryotic septins that compartmentalize the cytokinesis of budding yeast and the bacterial dynamin homologs required for high fidelity cell division during sporulation of *Streptomyces coelicolor* (20–23, 48).

## ACKNOWLEDGEMENTS

We thank Felix Dempwolff, David Kysela, Sampriti Mukherjee, and Stephen Olney for technical support. We thank Ethan Garner for the ZapA-mNeongreen and mNeongreen-FtsZ fusion, and Xindan Wang for the *sacA::hbsu-mCherry cat* fusion, and the IU Nanoscale Characterization Facility for use of its instruments. The work was supported in part by FAPESP grant 16/05203-5, NIH grant R35 GM131783 to DBK and NIH grant R01 GM113172 to SCJ.

---

**Figure S1. Diagram of the fluid layer of the microfluidic device.** Black lines represent microfluidic channels; open circles are media/reagent input/output reservoirs; closed circles designate the location of peristaltic valves; and three valves in series are used as a peristaltic pump. Location of the microchannel array imaged by fluorescence microscopy is indicated by the red box. The same device was used previously in (32).

**Figure S2. Quantitative analysis of micrographs.** A) Histogram of total FtsZ fluorescence intensity in individual cells for wild type (gray) and a *noc* mutant (orange). For each frame, a line scan through the longitudinal axis of the cell was performed, and total FtsZ fluorescence intensity was measured by integrating the area under the line scan. B) Histogram of individual cell lengths for wild type (gray) and a *noc* mutant (orange). Data in panels A and B were taken from 13,000 wild type (DK5133) for > 420,000 measurements and 5000 *noc* mutant (DK5820) for > 170,000 measurements. C-H) 100 cells each of wild type (DK5133, gray) and a *noc* mutant (DK5820, orange) were manually tracked and measured through a cell division cycle to generate the data for each panel. C) Histogram of the Z-ring appearance period defined as the time between the appearance of one Z-ring and the next Z-ring. Note, in the case of the *noc* mutant, the Z-ring that will form the future division site first appears by splitting from the pre-existent Z-ring. Thus, for the *noc* mutant, the appearance period was defined as the time at which the Z-ring visibly splits from its progenitor. D) Histogram of the Z-ring persistence period defined by the time between the appearance of a Z-ring and the disappearance of that Z-ring. E) Histogram of the Z-ring polar duration defined as the time between a septation event and the disappearance of the Z-ring responsible for that septation event. F) Histogram of the Z-ring medial delay defined as the time between a septation event and the appearance of a Z-ring that will eventually give rise to the next medial division event. Note that the Z-ring medial delay of the *noc* mutant was negative on average because the medial Z-ring that would eventually promote cell division was formed by splitting from the primary focus in the preceding generation. G) Histogram of the Z-ring maturation period defined as the time between Z-ring appearance and when that Z-ring achieved peak local intensity. H) Histogram of the cytokinetic period defined as the time between Z-ring appearance and septation directed by that Z-ring. I) Histogram of the segregation time of individual chromosomes of wild type (DK5712, gray) and a *noc* mutant (DK6372, orange) measured by microscopic analysis. Chromosome segregation events were defined by a local 40% decrease in HBsu-mCherry fluorescence intensity below a threshold value. > 300 division events were counted per dataset. J) Histogram of ZapA density in individual cells measured by dividing the total ZapA-mNeongreen fluorescence intensity by cell length for wild type (gray) and a *noc* mutant (orange).

**Figure S3: In the *noc* mutants, ZapA does not migrate to the future division size with FtsZ.** Kymograph analysis of wild type (DK8138, A) and a *noc* mutant (DK8172, B). Cytoplasmic mCherry signal false colored red and ZapA-mNeongreen intensity false colored cyan. To assemble a kymograph, a single microfluidic channel is imaged at 2 min intervals. All images reproduced at the same magnification.

**Movie S1: Wild type growth in microfluidic channels with fluorescent FtsZ.** Constitutive cytoplasmic mCherry false colored red and mNeongreen-FtsZ false colored green. Strain DK5133. Movies are 100 frames over 200 min at a rate of 1 frame/2 min.

**Movie S2: *noc* mutant growth in microfluidic channels with fluorescent FtsZ.** Constitutive cytoplasmic mCherry false colored red and mNeongreen-FtsZ false colored green. Strain DK5820. Movies are 100 frames over 200 min at a rate of 1 frame/2 min.

**Movie S3: Wild type chromosome segregation in microfluidic channels with fluorescent**

**FtsZ.** HBsu-mCherry false colored purple and mNeongreen-FtsZ false colored green. Strain DK5712. Movies are 100 frames over 200 min at a rate of 1 frame/2 min.

**Movie S4: *noc* mutant chromosome segregation in microfluidic channels with fluorescent FtsZ.** HBsu-mCherry false colored purple and mNeongreen-FtsZ false colored green. Strain DK6372. Movies are 100 frames over 200 min at a rate of 1 frame/2 min.

**Movie S5: Wild type growth in microfluidic channels with fluorescent ZapA.** Constitutive cytoplasmic mCherry false colored red and ZapA-mNeongreen-false colored cyan. Strain DK8138. Movies are 100 frames over 200 min at a rate of 1 frame/2 min.

**Movie S6: *noc* mutant growth in microfluidic channels with fluorescent ZapA.** Constitutive cytoplasmic mCherry false colored red and ZapA-mNeongreen-false colored cyan. Strain DK8172. Movies are 100 frames over 200 min at a rate of 1 frame/2 min.

**Movie S7: *noc* mutant containing an inducible construct of ZapA and fluorescent FtsZ grown in the microfluidic channel in the presence of 0.01 mM IPTG.** Cell body (*P_sigA_-mRFP mars*) false colored red and mNeongreen-FtsZ false colored green. Strain DK8144. Movies are x100 frames over 200 min at a rate of 1 frame/2 min.

